# Criticality explains structure-function relationships in the human brain

**DOI:** 10.1101/2024.07.15.603226

**Authors:** Marianna Angiolelli, Silvia Scarpetta, Pierpaolo Sorrentino, Emahnuel Troisi Lopez, Mario Quarantelli, Carmine Granata, Giuseppe Sorrentino, Vincenzo Palmieri, Giovanni Messuti, Mattia Stefano, Simonetta Filippi, Christian Cherubini, Alessandro Loppini, Letizia Chiodo

**Affiliations:** Department of Engineering, Università Campus Bio-Medico di Roma, Rome, Italy; Department of Physics “E.R.Caianiello”, University of Salerno, Fisciano, Italy; INFN sez. Napoli Gr. Coll. Salerno, Italy; Aix Marseille Univ, INSERM, INS, Inst Neurosci Syst, Marseille, France; Department of Biomedical Sciences, University of Sassari, Sassari, Italy; Department of Medical, Motor and Wellness Sciences, University of Naples “Parthenope”, Naples, Italy; Institute of Diagnosis and Treatment Hermitage Capodimonte, Naples, Italy; Institute of Applied Sciences and Intelligent Systems of National Research Council, Pozzuoli, Italy; Dipartimento di Scienze fisiche e tecnologie della materia del Italian National Research Council (CNR-DSFTM), Piazzale Aldo Moro 7, Rome, Italy; Istituto Nazionale di Ottica del Italian National Research Council (CNR-INO), Largo Enrico Fermi, 6, Florence, Italy; Department of Science and Technology for Sustainable Development and One Health, Università Campus Bio-Medico di Roma, Rome, Italy; Department of Medicine and Surgery, Università Campus Bio-Medico di Roma, Rome, Italy

**Keywords:** brain criticality, cortical dynamics, tractography, modular neural network, plasticity, magnetoencelography data, connectome

## Abstract

Healthy brain exhibits a rich dynamical repertoire, with flexible spatiotemporal patterns replays on both microscopic and macroscopic scales. How do fixed structural connections yield a diverse range of dynamic patterns in spontaneous brain activity? We hypothesize that the observed relationship between empirical structure and functional patterns is best explained when the microscopic neuronal dynamics is close to a critical regime. Using a modular Spiking Neuronal Network model based on empirical connectomes, we posit that multiple stored functional patterns can transiently reoccur when the system operates near a critical regime, generating realistic brain dynamics and structural-functional relationships. The connections in the model are chosen as to force the network to learn and propagate suited modular spatiotemporal patterns. To test our hypothesis, we employ magnetoencephalography and tractography data from five healthy individuals. We show that the critical regime of the model is able to generate realistic features, and demonstrate the relevance of near-critical regimes for physiological brain activity.

## 1 Introduction

The human brain dwells in a highly responsive state [1] where correct behavioural responses to multiple stimuli can be promptly selected among a large stored repertoire. This responsive state is mirrored by whole brain activities which constantly reconfigure themselves, generating a rich dynamical repertoire, despite the largely static anatomical connections [2, 3]. Such large dynamical repertoire manifests as numerous configurations of brain neurophysiological activities over time, as typically observed in healthy human brains. The functional repertoire shrinks in neurodegenerative diseases [3, 4], proportionally to clinical impairment, which suggests its possible use as a marker of the spreading of pathophysiological processes. Topology of the anatomical connections (structural connectome) alone cannot account for the fast, flexible dynamics observed in health. What other “ingredients” are needed for the brain to express a healthy functional repertoire? We hypothesize that the microscopic dynamics should operate in, or near, a critical regime for the functional repertoire to be effectively explored and for realistic flexible dynamics to emerge. A long-held theory posits that the brain self-organizes into a critical regime near the edge of a dynamical phase transition [5–9], where the balance between global inhibition and excitation plays a crucial role [10–12], and possibly leading to functional optimality [10, 13–19]. It has been proposed that the brain experiences both continuous phase transitions with scale-free avalanches, and discontinuous transitions, which are beneficial for self-sustained replay activity and memory functioning [20–22]. The roles of topology and learning have also been investigated [11, 23–27], with some notable insights on the role of modularity [28, 29].

To test our hypothesis, we build a modular Spiking Neuronal Network model to simulate collective neuronal dynamics emerging from the microscopic activities. Each module in this model consists of a group of Leaky Integrate and Fire (LIF) neurons, each representing a different cortical region. The network is trained using as starting point the structural connections on a probabilistic basis, for a series of modular spatiotemporal patterns through a learning rule [30] based on spike-timing-dependent plasticity (STDP) [31, 32], in this way establishing connections among neurons. A pattern is defined at the macroscopic level as a sequence of module activations, and at the mesoscopic level as a sequence of spikes. Activations propagate from one module to another based on a probabilistic distribution linked to the number of white-matter fibers, for long-range connections, and to an exponential decay rule, for close-by regions [33, 34]. As a result, the information (i.e. the learned spatiotemporal patterns) is encoded spatiotemporally, both as a precise sequence of neurons as well as a trajectory of activated modules.

After the learning phase, we focus on the ability of the model to spontaneously retrieve a maximally large number of previously stored patterns, which manifests as numerous transients, intermittently re-occurring events, as a function of varying underlying dynamical regimes. The dynamics of the model are characterized in terms of either hysteresis and bistability (generally associated with dynamics in a first-order phase transition) or scale-free avalanches (regarded as evidence of second-order phase transitions). To quantify the ability of different dynamical regimes to recover the previously stored large-scale functional motives and to generate realistic large-scale dynamics, we leverage large-scale bursts. They are observed in large-scale neurophysiological data in healthy brains [7] and their reconfigurations can be used to measure the functional repertoire in healthy and diseased brains [3, 35].

A growing body of experimental research, based on diverse recording methods at various temporal and spatial scales (e.g. fMRI, optical imaging, and magneto/electrical recordings), has consistently shown that spontaneous intrinsic neural activity exhibits correlations among neurons, cortical columns, and across extensive neuroanatomical systems [7, 36–41]. These correlations in spontaneous activities lead to the emergence of wave-like propagation of activities that are observable both at the mesoscopic and the macroscopic level.

The predictions of our model, in terms of the relationship between the structural connectome and the emerging dynamics, are compared to the empirical relationships observed between the structural bundles and the brain activities, as measured in source-reconstructed magnetoencephalographic (MEG) data acquired in five young healthy adults.

Under our hypothesis, we expect patterns to effectively spread across large scales, without runaway activities, near a critical regime. As such, the empirical correlation measured with MEG should be approached by the synthetic data when the system is operating at criticality.

## 2 Results

### Characterization of the model dynamics and hysteresis

A schematic illustration of the process of maturation of the model connectivity matrix during the learning stage is shown in (Fig. 1) (see also Methods for details). To forge the connectivity matrix, we use an STDP-based plasticity rule driven by modular patterns of activity extracted using the subject-specific anatomical information on structural connectivity. These modular patterns of activities are encoded in the network during the learning stage when a “teaching signal” inducing cortical plasticity is provided [36,42]. In this sense, this accommodates in our model an Hebbian learning rule. In particular, we use a combination of tractography (for far-apart regions) and the geometric distance among the centroids of the regions (for close-by regions, see Methods) to extract sequences of active modules in each pattern.

**Figure 1:**
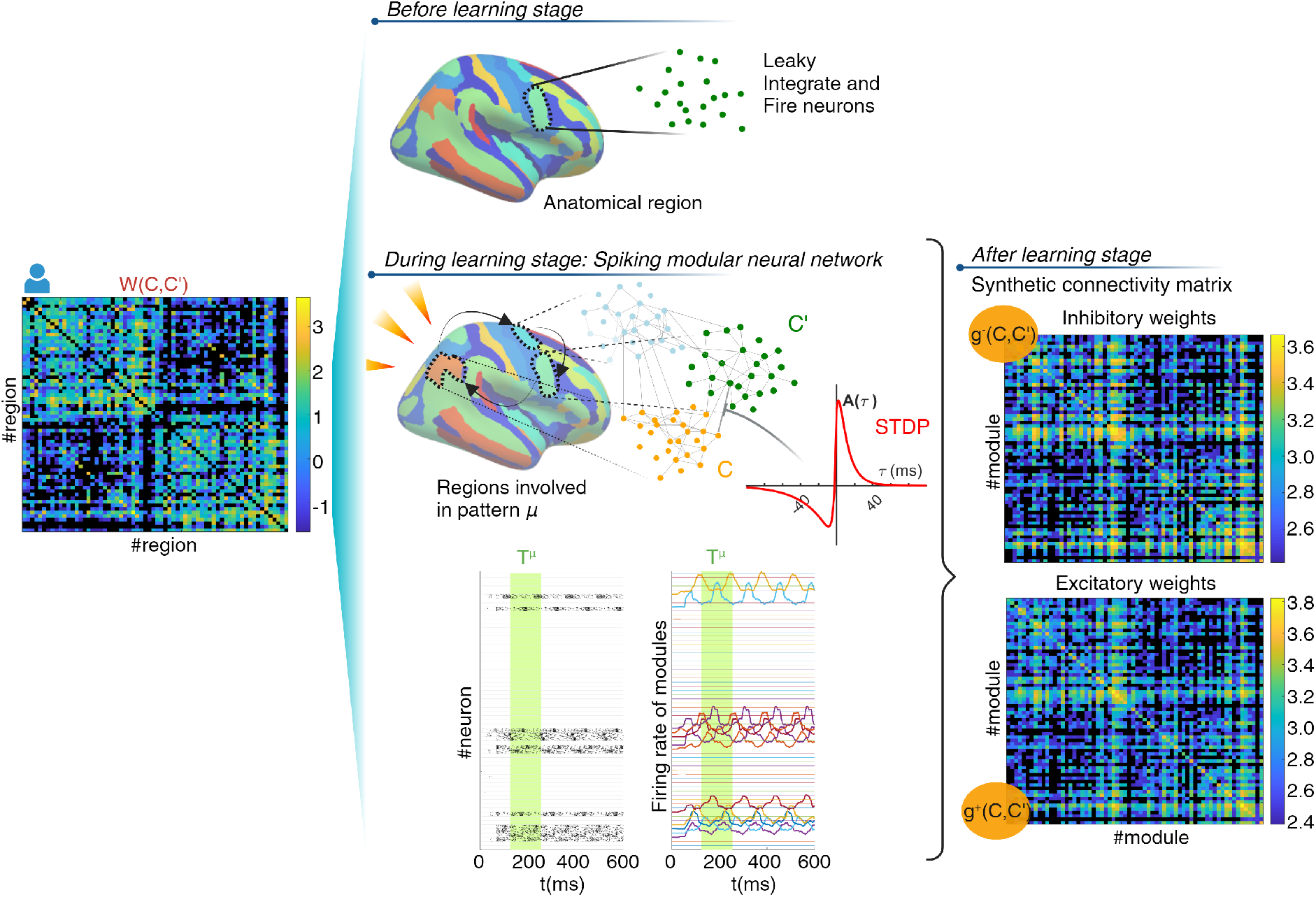
Schematic representation of the network. Left panel, empirical structural connectivity matrix *W* (*C, C*^*′*^) between the M = 66 cortical regions of interest (ROI). It includes either the number of white tracts or, for nearby regions, a term decaying exponentially with the Euclidean distance. The color scale ranges from blue to yellow, representing the logarithm of *W* (*C, C*^*′*^), while entries where *W* (*C, C*^*′*^) = 0 are displayed in black. Central panel, overview of the construction of the model. From the top: starting from disconnected neurons in each ROI (“module” in the model formalism), during the learning phase, the neural network is forced to learn a set of specific modular dynamic patterns through information extracted from the empirical connectivity matrix *W* (*C, C*^*′*^). Each pattern consists of a distinct spiking activity that spreads among particular neurons in specific modules, while others remain inactive. The connection weights between neurons are defined according to a learning rule based on spike-timing dependent plasticity (STDP), which is used to store the extracted patterns. At the bottom: an example of the rasterplot of one modular pattern used to train the network, whose period is denoted by a green rectangle; and the firing rates of various regions overlap. Right panel, macroscopic connectivity matrices *J*_*ij*_ between neurons after the learning stage, depicting the macroscopic excitatory (*g*^+^(*C, C*^*′*^)) and inhibitory connectivity (*g*^+^(*C, C*^*′*^)), respectively. The color spectrum from blue to yellow indicates the logarithmic strength, with black indicating null values. Pictures created with BioRender.com

After learning, the network converges on connections weights that, given the appropriate parameters, can give rise to multistable neural dynamics (i.e. with multiple attractors). The spiking neural network can retrieve previously learned patterns of correlated activities as waves of activations traveling across brain regions at the macroscopic level and as precise spatiotemporal dynamics of the spiking neurons at the microscopic scale. In fact, by modulating the strength of the weights and the global inhibition, one can have the model dynamics go from a quiescent state to a high correlated state of self-sustained replay. In the sustained state, one of the stored patterns is periodically recalled with high fidelity, i.e. the system replays precisely the sequence of regions [*C*^1^, …, *C*^*G*^] and the sequence of neurons inside each region. At low value of global inhibition *I*_0_, these two dynamical regimes are separated by a hysteretic region characterized by bistable dynamics where there are either quiescent or sustained replay regimes (Fig. 2, top and middle rows) depending on the history of the system. To assess the accuracy of the replays, we introduce an overlap parameter *q*^*µ*^ which quantifies the similarity of the spontaneous spiking activity with each pattern stored in memory *µ*, for *µ* = 1 … *P*. Then, we use the maximum over the *P* overlaps *q*^*µ*^ as an order parameter *Q* to characterize the overall dynamics of the system. A high value of *Q* indicates a strongly correlated activity, similar to one of the stored modular spatiotemporal patterns. We report in Fig. 2 the spontaneous firing rate and the order parameter during a hysteresis loop, where *E*_0_ is increased from 0 to 10 over 20 s and then decreased back to 0 in the same duration while keeping constant *I*_0_ and the noise intensity. Simulating the adiabatic changes (slowly and constantly changing the parameters) reveals the presence of hysteresis, for the low levels of the global inhibition *I*_0_ (Fig. 2, top and middle row). The discontinuity in the firing rate and in the order parameter, and the presence of hysteresis suggest a (dynamical) first-order phase transition. The area of the hysteretic loop gradually shrinks as the global inhibition increases, up to the critical value of *I*_0_ at which the hysteretic cycle is closed, the transition becomes continuous (Fig. 2, from upper to lower rows), and the system operates with critical-like properties. When the dynamics operate near a second-order phase transition, the stored traveling-wave patterns are no longer persistent attractors, and the spontaneous dynamics shows a variety of transient short replays with high fluctuations of activities and a high number of reconfigurations.

**Figure 2:**
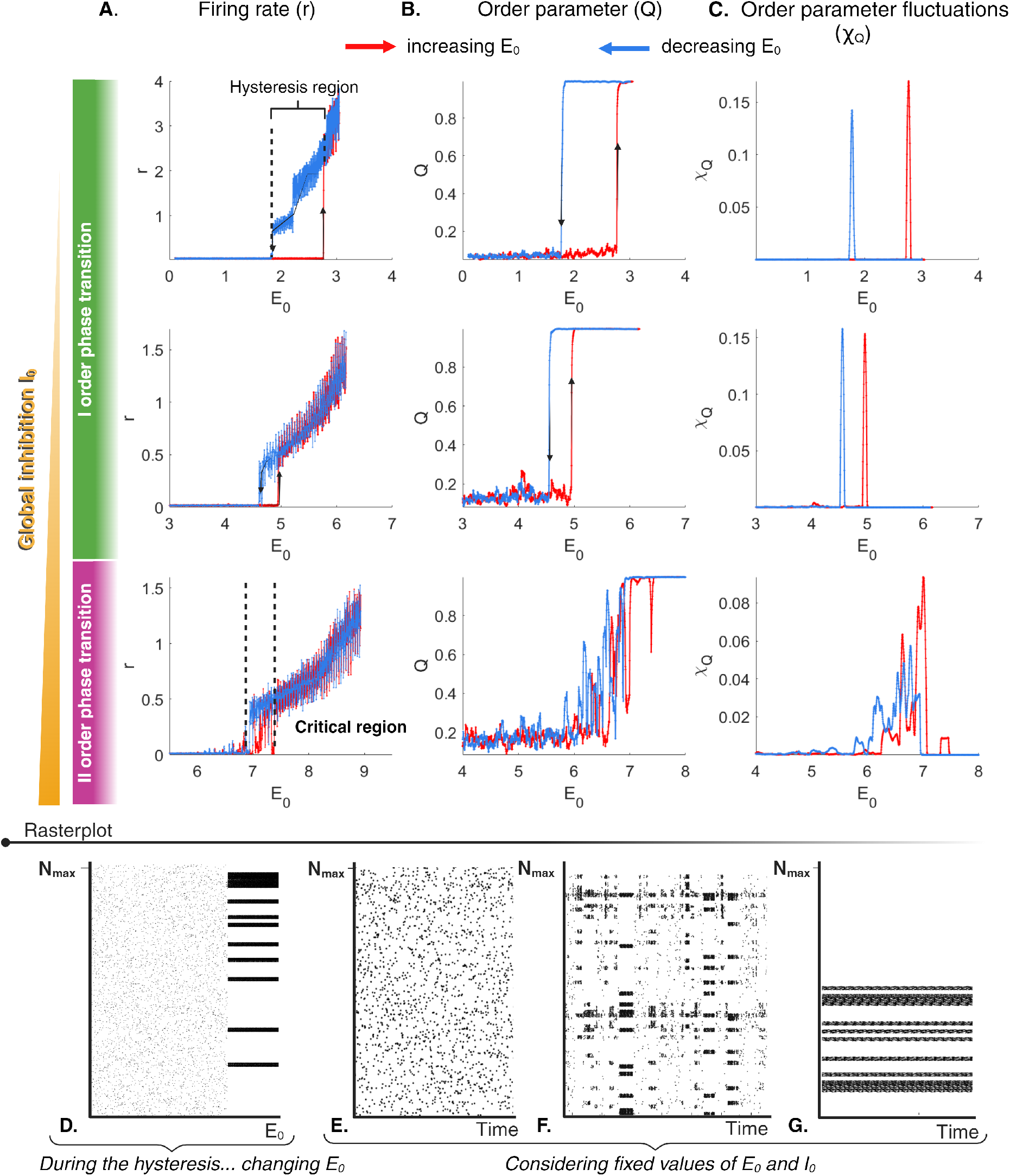
From first-to second-order phase transitions. The spontaneous dynamics of the network as a function of the excitatory connectivity parameter *E*_0_ during a hysteresis loop. Panels A-C: firing rate of the entire population over 1 ms interval (panel A), dynamics of the order parameter (evaluated in a sliding window of 200ms) capturing transitions in the network activity (panel B), fluctuations of the order parameter (panel C). The dynamics are explored while systematically varying the parameter *E*_0_ (shown on the *x*-axis of each plot), given a constant noise level (*α* = 0.5, *ρ* = 1*ms*^*−*1^) and different fixed levels of global inhibition (*I*_0_ = 0.3, 0.8, 1.1 from top to bottom rows). The red and blue lines in A-C refer, respectively, to the increases or decreases of excitation during the simulation. Panel D: At lower global inhibition *I*_0_, the network shows a hysteretic discontinuous phase transition characterized by a sudden shift from low to high firing rates marked by the replay of one stored pattern and a jump in the order parameter from 0 (indicating uncorrelated activity) to 1 (indicating a perfectly retrieved pattern) as shown in the rasterplot. As global inhibition increases, the area of the hysteresis loop decreases, and the transitions become smoother, characteristic of a second-order phase transition. This is evident from the broad peak in the order parameter fluctuations, indicating a critical regime. Panel D shows network activity while *E*_0_ increases for *I*_0_ = 0.3 where the transition to the self-sustained collective state is discontinuous. Panels E-G: dynamics for *E*_0_ = 5, *I*_0_ = 1.1, *E*_0_ = 6.9, *I*_0_ = 1.1 and *E*_0_ = 7.9, *I*_0_ = 1.1 respectively, which are an example of subcritical, critical and supercritical regime, respectively.

While in the sustained replay regime (i.e. the supercritical region) the replays are exact replicas of one stored modular pattern, at the boundary between the uncorrelated and the strongly correlated regime, more patterns are briefly replayed, but each of them with lower accuracy.

In the parameter region where the network exhibits first-order phase transitions, the fluctuations of the order parameter show very sharp peaks at the edges of the hysteretic cycle, which trivially reflects the discontinuity of the order parameter. However, by increasing the global inhibition, the critical region is approached and the peak of the order parameter fluctuations (*χ*_*Q*_) becomes wider. To better characterize this regime, it is useful to analyze the temporal dynamics with static control parameters (with fixed values *E*_0_ and *I*_0_). In the limit of static dynamics averages (which can be understood as an infinitely slow - that is, static-adiabatic change), the bistability region collapses into a line of a discontinuous transition. Fig. 3 shows the mean value of the order parameter (computed over 200ms long windows and averaged over the total simulation time of 20.000ms), for static conditions runs for different combinations of fixed *E*_0_ and *I*_0_. A supercritical region appears for large *E*_0_ and low *I*_0_, where the order parameter is close to 1, which means that the spontaneous collective dynamics shows self-sustained replays of one of the stored patterns. For large *I*_0_ and low *E*_0_ values, there is a subcritical region where the order parameter is near zero. Here firing rate is low. Between these two regimes, we can identify a critical region (in the upper right triangular area) where the fluctuations of the order parameter (over time, for fixed control parameters) are very large (see Fig. 3, panel B). To demonstrate that these large fluctuations of the order parameter over time can be traced back to the nature (i.e. first or second order) of the underlying phase transition we overimpose to the (*E*_0_, *I*_0_) plane the points where the dynamics, observed for each constant *I*_0_ while slowly changing *E*_0_, switches from a low (*Q <* 0.9) to a high value (*Q >* 0.9) of the order parameter. The resulting lines are called pseudo-spinodals. The lower pseudo-spinodal, dashed black lines in Fig. 3, shows the point where the dynamics goes from low to high values of the order parameter, and the upper pseudo-spinodal, continuous black line, shows the point where the dynamics goes from high to low values of the order parameter, during hysteresis loop (also shown in Fig. 2). Between the two pseudo-spinodals, there is a region of bistability. The two pseudo-spinodals collapse where the hysteresis loop disappears and, thence, a second order transition occurs.

**Figure 3:**
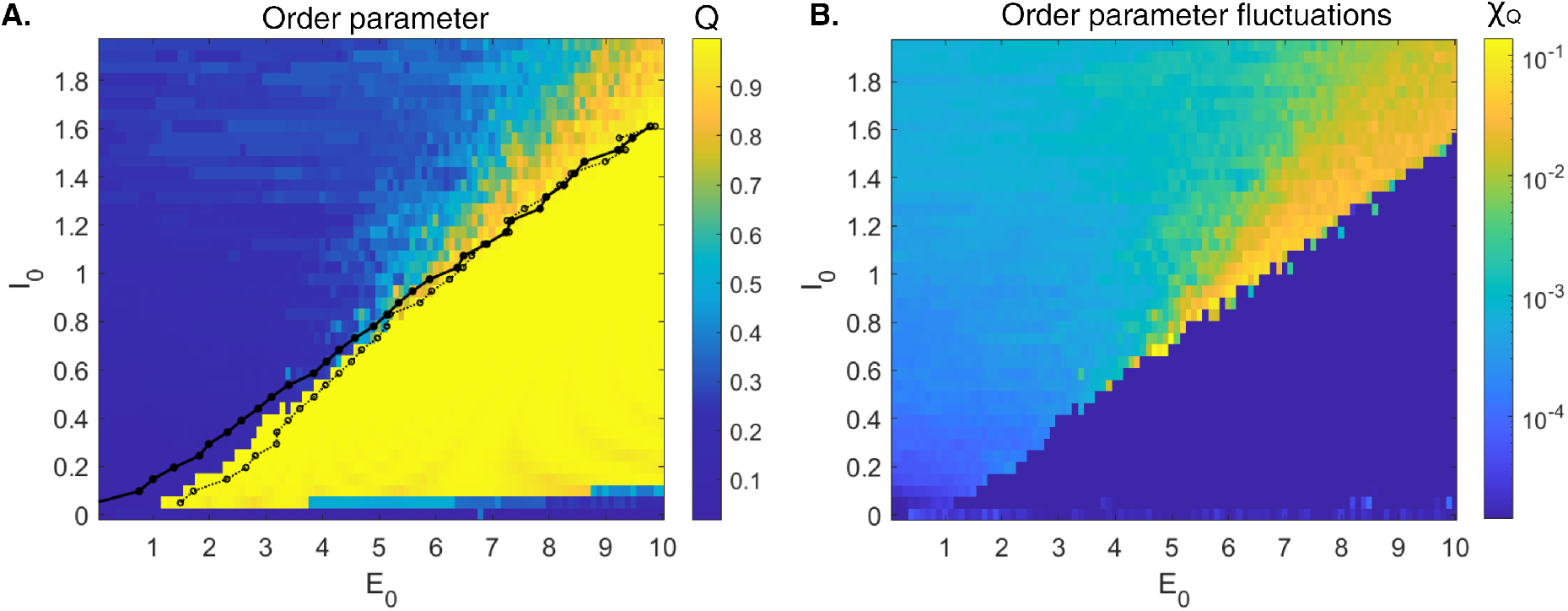
Order parameter and its fluctuations. The average of the order parameter over time for different fixed values of parameters *E*_0_ and inhibition *I*_0_ is displayed (Panel A) along with the fluctuations of the order parameter (Panel B). We observe a region of the parameter space where the order parameter is close to 1 and exhibits very low fluctuations. At lower values of *E*_0_, there is a regime with low values of the order parameter. Between these two regimes, an extended region opens up where the order parameter has an intermediate mean value and large fluctuations over time. In panel A we overimpose to the (*E*_0_, *I*_0_) plane the pseudo-spinodal line, that is the *E*_0_ values at which, during a hysteresis sweep, a transition from a self-sustained collective dynamics to a low-rate state occurs. The other (dashed) pseudo-spinodal line shows where the opposite transition (i.e. from low rate to self-sustained dynamics during increasing *E*_0_) occurs. Note that the two lines converge for high values of *I*_0_. This suggests a change in the system’s dynamic responses and state transition properties depending on global inhibition and noise level (*α*). Additionally, where the two pseudo-spinodal lines begin to overlap, the mean order parameter at stationary conditions shows a smooth change as a function of *E*_0_. In this area of the parameter space, the fluctuations of the order parameter reach their maximum, as shown in the right panel (using a logarithmic scale for the color bar). The large fluctuations of the order parameter observed in this extended region, near the edge of the transition, are a signature of critical behaviour in the network dynamics, emphasizing the interplay between excitation, inhibition, and noise in shaping the system’s dynamics.

We observe that the line of first-order transitions ends (the spinodal lines meet) at about *I*_0_ *≃* 1, *E*_0_ *≃* 6, where an extended critical region with large fluctuations of the order parameter opens up. This is reminiscent of the “Widom line” [43, 67] (the locus of stability minima and order-parameter fluctuation maxima). In this upper-right triangular region, one observes small transient collective replays of some stored modular patterns, leading to large fluctuations of the order parameter. In conclusion, our results demonstrate that transient collective replays and large fluctuations of the order parameter occur in this extended region between the two dynamical states.

We further characterize this critical region utilizing the Fano Factor and the coefficient of variation (CV) of the inter-spike intervals under static conditions, as a function of *I*_0_ and *E*_0_.

### Analysis of the critical region

In Fig. 4, the Fano factor (see eq.14) computed over a bin width δ_bin_ = 200*ms* (refer to Methods for the rationale behind using large time bins) and averaged over the last 5s of a stationary simulation, is shown as a function of *E*_0_, *I*_0_. There is an extended region where the Fano factor is large, corresponding to the regime with large order parameter fluctuations, as shown in Fig. 3.

**Figure 4:**
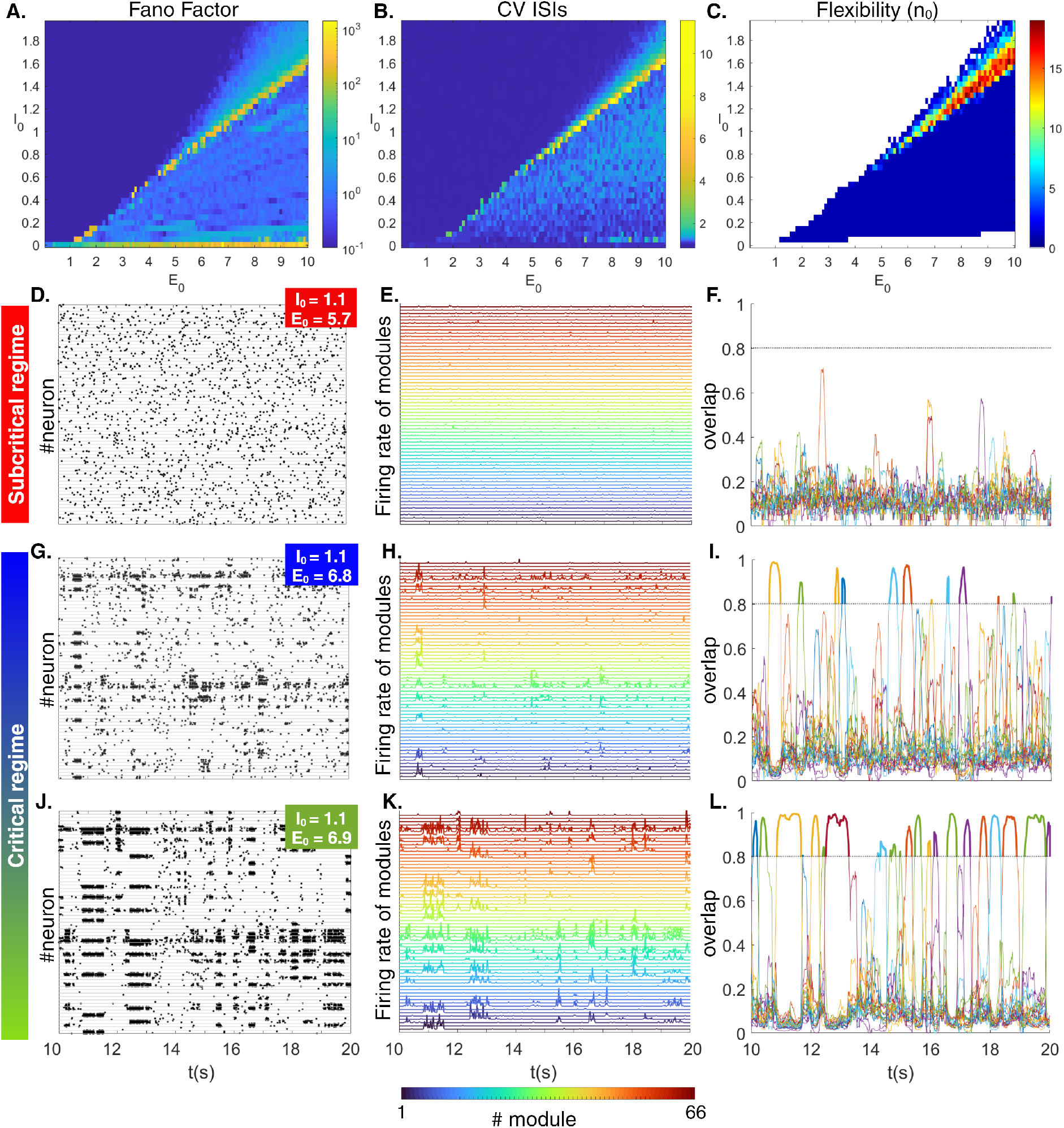
Characterization of critical region in simulated data. The three pictures at the top, show the Fano factor (panel A), the Coefficient of variations of Interspike intervals ISIs averaged over modules (panel B), and the flexibility, i.e. the number *n*_0_ of distinct reactivated patterns (panel C), as a function of the global inhibition *I*_0_ and strength of the structural connections *E*_0_. These measures are used to characterize the parameter space. We show the rasterplots, the firing rates of modules and the overlap as a function of time for one example of subcritical regime (panels D, E and F) (considering *I*_0_ = 1.1, *E*_0_ = 5.7), and two examples of critical regimes (panels G, H, I and J, K, L) (considering the values *I*_0_ = 1.1, *E*_0_ = 6.8 and *I*_0_ = 1.1, *E*_0_ = 6.9). In all the regimes, as shown in panels F, I, L a threshold of 0.8 (indicated by the dashed line) is used to highlight a period of high similarity between the spontaneous activity and the stored one.

Similarly, the CV of the interspike intervals among spikes of the modules (panel B of Fig. 4), averaged over modules, also shows high values in the region with large fluctuations of the order parameter. To summarize, we have characterized an extended area of the parameter space, reminiscent of a Widom line, by large fluctuations of the order parameter, a large Fano factor, and CV ISIs. This region appears near the edge of the transition between the persistent collective state and the low firing rate state.

Then, for each simulation at fixed parameters *E*_0_ and *I*_0_, we count the number *n*_0_ of distinct patterns that were transiently replayed with an overlap *q*_*µ*_ (measured over a time window of 200*ms*) greater than 0.8. The number *n*_0_ of replayed distinct patterns is zero in the area to the left of the parameter space (where inhibition is large) and is 1 in the regime of self-sustained persistent replay to the right (panel C of Fig. 4. Following our predictions, the extended critical region, characterized by significant fluctuations in the order parameter, also exhibits a higher number of transiently and intermittently replayed patterns. This is due to the coexistence of correlated and uncorrelated activity and the intermittent emergence and fading of stored patterns in this critical regime.

We report examples of the raster plots of the network activity and modules’ rate as a function of time in Fig. 4. Looking at the two lower raster plots, which refer to the dynamics operating in the critical dynamic regime (at *E*_0_ = 6.8, 6.9 and *I*_0_ = 1.1), a stochastic switching of states appears, whereby a modular pattern spontaneously emerges and quickly fades away, giving rise to a sequence of recalled patterns alternating with states of low rate. Such dynamics is also demonstrated in Fig. 4, panels *F, I, L*, where each coloured line refers to the overlap of the ongoing activity with a different stored pattern, as a function of time. It is easy to see that in the critical regime high fluctuations occur, with numerous patterns being alternatively recalled over time. Conversely, in the subcritical region (at *E*_0_ = 5.7 and *I*_0_ = 1.1), where the mean order parameter is close to zero, the system is unable to recall any pattern. Finally (not shown), in the supercritical regime the dynamics become entrained to one specific pattern. These stereotyped dynamics are less susceptible to external stimuli (and to noise) as compared to the critical regimes characterized by large fluctuations. To further explore the spatiotemporal structure of transient events, we examine the size and duration distributions of intermittent bursts of activity (Fig. 5), usually referred to as avalanches (see definition in Methods) at the same values of (*E*_0_, *I*_0_) used in Fig. 4, panels D-L. In the critical regime, size and durations of neuronal avalanche distributions are better described by power law, *P* (*S*) *∝ S*^*−α*^ and *P* (*T*) *∝ T*^*−β*^, respectively, than by exponential in the range shown in Fig. 5. Power laws indicate the absence of characteristic scales in the underlying dynamics and are therefore considered a hallmark of criticality. Interestingly, as we will see later, power law behavior is also consistent with MEG empirical data.

**Figure 5:**
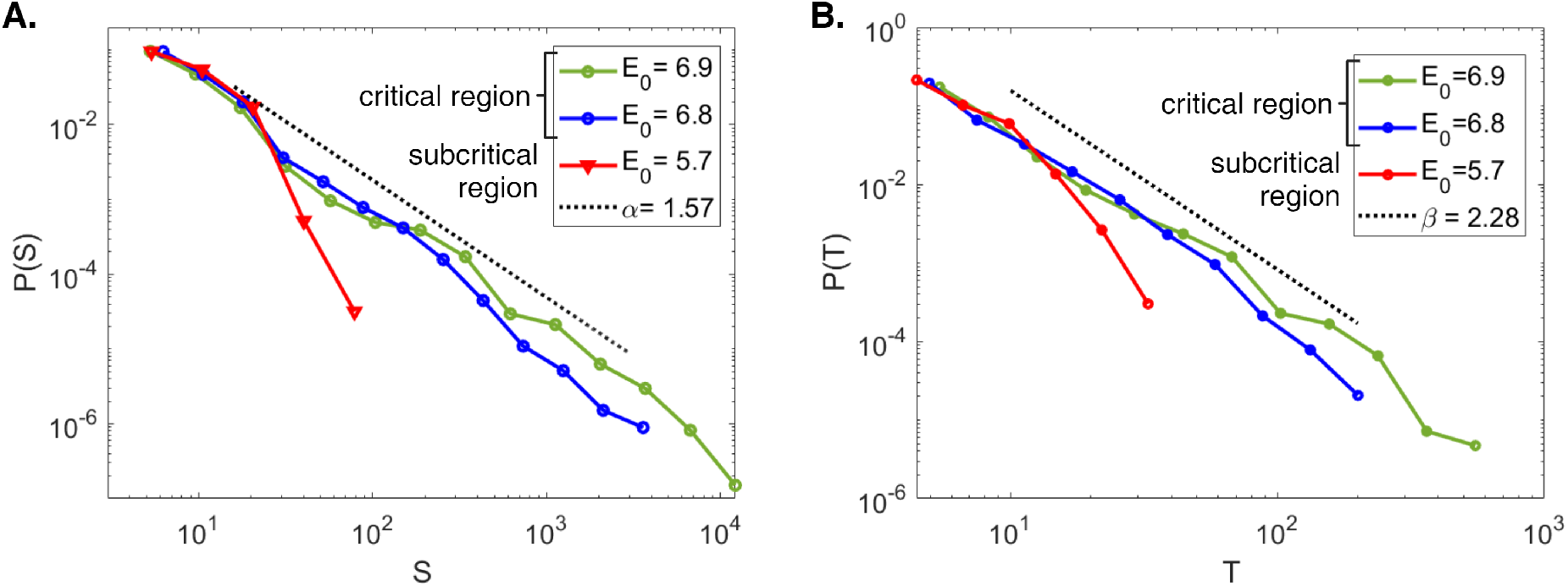
Neuronal avalanches in synthetic data. Panels A-B: distribution of the size and durations of the neuronal avalanches, respectively. For each panel, the values of *I*_0_ = 1.1 and *E*_0_ are used (see also 205 Fig. 4, panels D-L). The dashed line indicates the fit performed using the powerlaw python package, for *E*_0_ = 6.9 and *I*_0_ = 1.1, over the range of values indicated by the black dashed line. The two distributions follow a power law with exponent *α* = 1.57 ±0.04 ±0.06 (fit± error on the fit± SE) for the size and *β* = 2.28± 0.07± 0.2 (fit ± error on the fit± SE) for the durations. The Kolmogorov-Smirnov distance between the data and the fit is *D* = 0.15 for the size and *D* = 0.09 for the durations. The log-likelihood ratio between the power-law and the exponential fit is *R* = 77(*p <* 10^*−*5^) for the size and *R* = 63(*p <* 10^*−*6^) for the durations, indicating that data are more likely to follow a power law than an exponential distribution (see Methods for details of the fit).

We now set out to verify whether the features expected in the critical regime can be observed in the empirical data. To this end, we use MEG source-reconstructed resting-state data from the MEG scans of the same five healthy young adults for which the structural connectomes are available. For these analyses, we consider only cortical regions, given the better signal-to-noise ratio of more superficial sources.

Firstly, we check if fluctuations of the z-scored MEG signals deviate from a Gaussian fit (Fig. 6) to choose the threshold (*σ*) useful to define a region as active. In this way, an avalanche is defined as a continuous time interval during which at least one z-scored MEG signal exceeded the threshold, either positively or negatively. The distributions of size and duration of neuronal avalanches, pooled over all the subjects, are shown in panel C-D of Fig. 6. Using the powerlaw package [44] we test that the power-law fit is more likely than the exponential one in the interval shown by the black line (see Methods).

**Figure 6:**
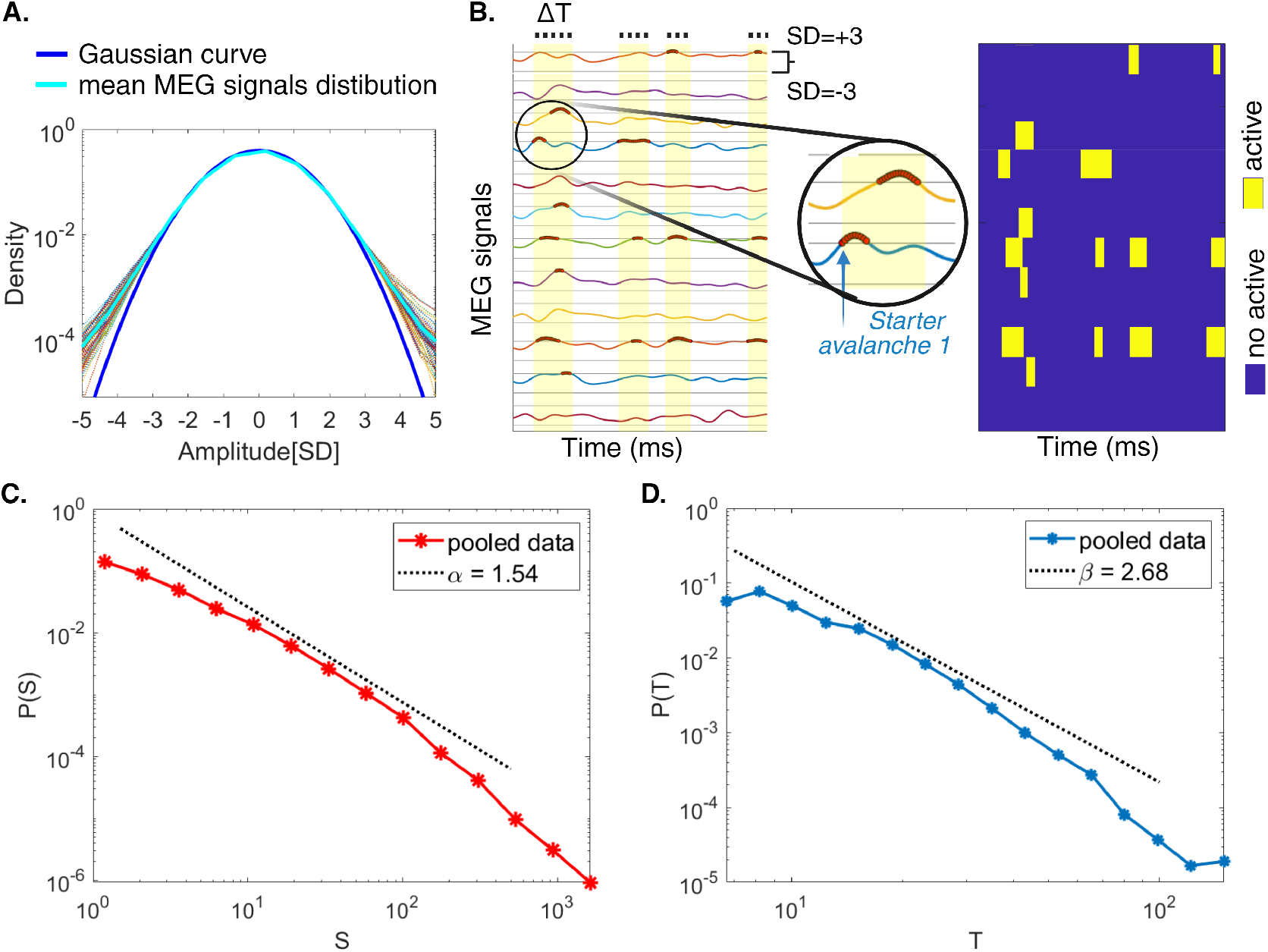
Empirical analysis of neuronal avalanches. Panel A: distribution of the z-transformed source-reconstructed magnetoencephalography (MEG) signals. The background (coloured) curves represent the signal amplitude distributions for all five subjects (based on all channels), with the light blue curve indicating the grand average across all subjects. The blue curve represents the best-fit Gaussian distribution for this grand average. Notably, the normalized data distributions start to deviate from the Gaussian fit around *θ* = ±3 SD. Panel B, left: example of MEG signals, where the signals crossing the threshold of ± 3 standard deviations (SD) are marked in red. For illustrative purposes, the initiating region of the first avalanche is indicated by the light blue arrows. Panel B, right: the raster plot shows in yellow the active regions. Panel C: avalanche size distribution across all subjects. It follows a power law with an exponent *α* = 1.54 ± 0.006 ± 0.05 (fit±error on the fit ± SE). Panel D: avalanche durations distribution across all subjects. It follows a power law with an exponent *β* = 2.68 ±0.02± 0.3 (fit ±error on the fit ±SE). In both cases, the fit has been done using the powerlaw python package over the range of values indicated by the black dashed line. The Kolmogorov-Smirnov distance between the data and the fit is *D* = 0.12 for the size and *D* = 0.10 for the durations. The log-likelihood ratio between the power law and the exponential fit is *R* = 191(*p <* 10^*−*57^) for size distribution and *R* = 86(*p <* 10^*−*2^) for durations, indicating that the data is more likely to follow a power law than an exponential distribution (see Methods for details of the fit).

### Correlation between brain anatomic structure and brain dynamics

As described in the previous paragraph, we provide extensive characterization of the model to demonstrate critical dynamics and its relation with the expected observable large-scale quantities. We now leverage the simultaneous availability of structural and functional data to test one more prediction derived from the *in silico* simulations. We test the hypothesis that the critical regime best justifies the observed correlation between the topology of the structural connectome and the corresponding topology of the spreading of neuronal avalanches. In other words, there is a well-known relationship between structural and functional data. We hypothesize that such a relationship is evident when the system operates in a critical regime. To this end, we compute the Spearman correlation (see eq. 16) between the degrees of the nodes (the number of streamlines connecting each node to any other node) obtained from the connectome, and the number of times each node could initiate a neuronal avalanche in synthetic data. In Fig. 7, is shown that the critical region is the one where the correlation between the nodal degree and the functional role of each region as “avalanche-starter” is higher.

**Figure 7:**
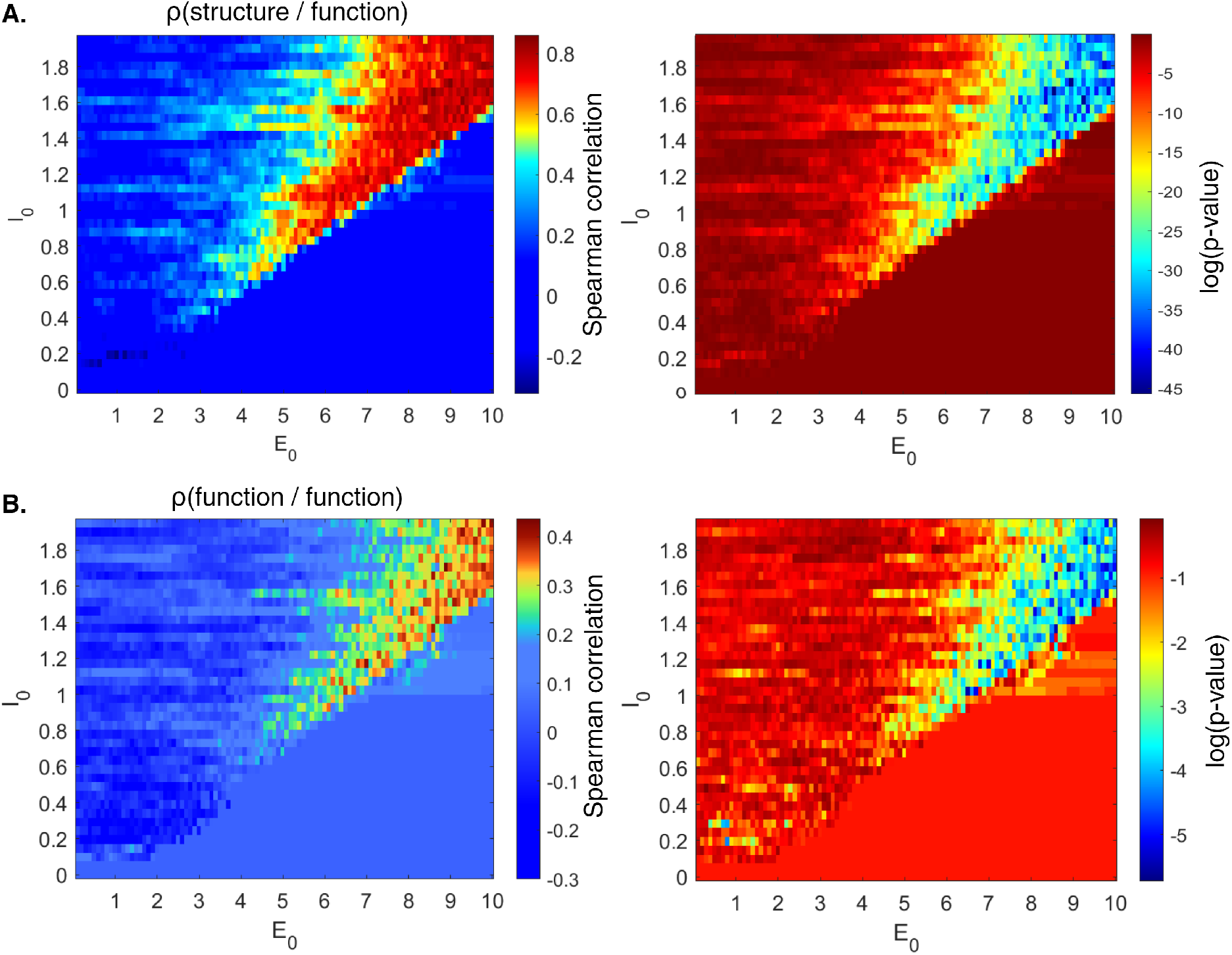
Structure-function relationship. Panel A, left: matrix of Spearman correlations *ρ* between the structural connectivity (empirical degrees of the nodes) and the simulated functional dynamics (number of times each node initiated a neuronal avalanche in synthetic data); panel A, right: associated p-values, both as a function of the global inhibition *I*_0_ and structural connections strength *E*_0_. Panel B, left and right: Spearman correlation between simulated and empirical functional dynamics, and associated p-values, respectively.

Finally, we compare directly synthetic and empirical functional data. Once again, the critical region is the one where the model best reproduces the topography of the “avalanche-starting” regions, as observed in Fig. 7. In conclusion, we show that, while the structural connection is not changed in multiple simulations, only if the model is set to a critical regime realistic large-scale dynamics can emerge.

## 3 Discussion

In this manuscript, we set out to provide an account of the ability of the brain to quickly and flexibly recruit multiple pieces of previously stored information (as represented by multiple specific patterns of activations), despite constant structural connections among brain regions, at a fast timescale. Empirically, one can observe that, despite the underlying structural connectivity constrains the patterns of activations across the brain, the dynamics are flexible, and a healthy brain can alternatively recruit numerous stored functional patterns (i.e. express a large functional repertoire).

As a result, while each burst of activation might recruit only a few regions, all the areas are recruited over time at some point (since the spatiotemporal dynamics of the bursts vary over time). Thence, bursts of activations tend to spread across all the structural tracts, whereby the observed structural-functional correlation emerges [68].

In detail, we hypothesize that the structure-function relationship is best understood as emerging from microscale and mesoscale dynamics operating in the critical edge of a second-order phase transition. The idea that some features of resting state dynamics can be captured only when models are tuned to criticality was also investigated in human fMRI experiments using the Greenberg-Hastings model [40].

To test this hypothesis, we use a Leaky Integrate and Fire model embedded into a multiscale network. That is, neurons are allowed to communicate either by proximity (for close-by neurons, according to an exponential decay rule), representing the microscale, or over the topology imposed by the large bundles of white matter tracts (as measured by the structural connectome), to accommodate the mesoscale.

The model exhibits two distinct dynamical regimes: an uncorrelated low-rate state and a strongly correlated state characterized by a high-order parameter. These regimes are separated by either a first-order or second-order phase transition, depending on the global inhibition strength. When the hysteresis loop narrows, a continuous phase transition occurs, creating an extended region at the boundary between the two states characterized by high order parameter fluctuations. This region is marked by numerous transients and intermittently recurring motif events. We show in the model the presence of a supercritical regime where the system can operate as an associative memory. In this regime, the self-sustained recruitment of a single activation pattern (from among many stored) serves as a persistent attractor in the dynamics, entraining the whole system. Hence, in a supercritical regime, the system loses the ability to recall more patterns spontaneously. In other words, the dominance of the persistent pattern restricts the system’s ability to transition to other states. The network, in essence, becomes confined to a narrow range of responses and can no longer produce dynamic and adaptable behavior. These findings talk to previous literature where it was shown that losing the flexibility of the dynamics during the resting state occurs in pathological conditions such as Parkinson’s disease and amyotrophic lateral sclerosis [3,4]. Furthermore, the flexibility of the brain dynamics is also regulated by physiological factors. For example, the flexibility of the dynamics during resting state varies as a function of the menstrual cycle, with the pre-ovulatory phase invariably showing maximized flexibility [45]. Typically, these findings are understood as healthy brains achieving readiness to numerous stimuli by being constantly partially “trying” multiple strategies. A tennis player jumping left and right (oscillating between two states) to be as ready as possible for multiple potential serves from the opponent represents an analogy [1]. As such, the dynamics we observed in supercritical regimes would not be ideal for constantly recalling multiple patterns. Conversely, a large sensibility to stimuli is expected in the critical regime. Our model shows that the system, in the critical regime, spontaneously fluctuates among quiescence and short-lived collective modular patterns (i.e. attractors), showing remarkable flexibility and adaptability. This is in line with the observation [46] that large-scale networks in resting-state MEG can be well described by repeated visits to short-lived transient brain states, defined as patterns of network activity, spatially and spectrally distinct, across a set of regions which span the whole brain. Notably, also experimental ongoing neuronal avalanches are composed of diverse repeating spatial patterns [47–49]. Moreover, recently it has been examined the role of fast GABA-receptor mediated synaptic inhibition, showing that the diversity and selectivity for families of ongoing avalanche patterns depend on inhibitory signalling [50].

Interpreting the presence of multiple replays as the implementation of memory mechanisms, our results can be framed within previous studies showing that cortical replay signals can occur at the whole-brain level [51, 52], and that spontaneous brain activity, measured with resting state fMRI, is correlated among regions that are co-activated by behavioural tasks [51]. Successful delayed memory recall was associated not only with increased hippocampal–cortical interactions but also with changes in the cortico-cortical patterns of co-activations [53].

In this work, we implement a model to study how a healthy brain operates near a critical point, including the cross-scale modularity of brain networks. The correlation between the critical region and the flexibility of the emergent dynamics is apparent when observing the system at multiple scales, both at the spike level and when the dynamics are translated to modules (i.e. to the higher organizational level). In other words, the behavior at the mesoscale mirrors the dynamics and flexibility observed at the microscopic level (spikes).

Both theoretical and empirical arguments show that modularity might be a parsimonious and cost-effective organizational principle of brain architecture. From a theoretical standpoint, sparsely interconnected modules allow faster adaptation or evolution of the system in response to changing environmental conditions. Of note, the formation of modules is typically observed in dynamical systems where network structure and function coevolve [54]. From an empirical standpoint, modularity in human anatomical networks has been shown [55].

When tuned in a critical regime, our model reproduces, in terms of neuronal avalanches, similar dynamics to the one observed in the empirical data. This aligns with signatures of criticality previously described in large-scale neurophysiological data in humans [7]. Furthermore, the use of the coefficient of variation of interspike intervals (CV of the ISIs) helps our understanding of the variability in neuronal firing. The CV of the ISIs focuses on the timing between individual spikes and is complementary to the Fano factor, which looks at the variability of the firing rate (offering a more direct link to the mesoscale dynamics). Altogether, these measures provide a multi-scale view of neural dynamics, from individual neuron behavior to collective network activities.

We then analyze the relationship between function and structure in the brain. Previous evidence showed that the spreading of neuronal avalanches in the human brain is constrained by the structural connectome over the large scale. In our study, we show that the empirical structure-function relationship is best explained by assuming that the underlying dynamics are operating in a critical regime. In other words, our findings suggest that the critical state allows the brain to achieve maximally flexible dynamics despite operating on a static structural scaffold. This provides a mechanistic link between the emergent properties of the brain networks and the underlying anatomical connectivity. In fact, our results suggest that the structural architecture of the brain facilitates its complex functions, with critical dynamics acting as a bridge that enables the translation of structural connectivity into optimal functional outcomes. Consequently, our research offers a comprehensive framework for understanding how the brain’s structure supports its diverse cognitive and processing capabilities, highlighting the importance of critical states in achieving optimal functional efficiency.

## 4 Methods

### The Modular Spiking Neural Model

We implement a modular network, where each module corresponds to one of M=66 cortical regions of interest (using the Desikan-Killiany-Tourville DKT atlas [56]). This design is grounded in the concept that the brain’s network is organized into distinct regions or modules. Within each module, neurons typically exhibit stronger connectivity among themselves compared to their connections with neurons in other modules.

Each of the *M* modules consists of an ensemble of *Z* = 200 Leaky Integrate and Fire (LIF) neurons, for a total of *N* = *MZ* = 13200 neurons in the network.

The membrane potential *V*_*j*_(*t*) of the *j − th* neuron, *j* = 1, .., *N* follows the equation

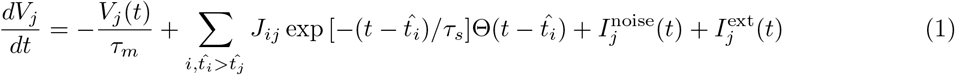

where the membrane time is *τ*_*m*_ = 10*ms* and the synapse time is *τ*_*s*_ = 5*ms* [57, 58]. When the potential reaches the threshold *θ* (conventionally set to *θ* = 1), the neuron fires and the potential is reset to the resting value *V*_*j*_ = 0. Θ is the Heaviside function, 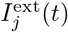 is the external input, taken zero when we study spontaneous dynamics. The noise term 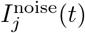,modelling spontaneous neurotransmitter release and other stochastic sources, is defined as:

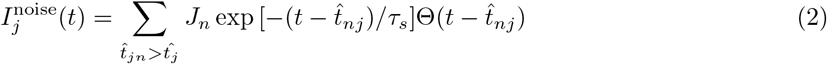

where *J*_*n*_ is extracted from a Gaussian distribution with mean equal to 0 and standard deviation *α*, and the times 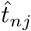 are extracted for each neuron *j* from a Poissonian distribution with a rate *ρ* = 1.

The connections *J*_*ij*_ between neurons are built during the learning stage by storing modular patterns of activity, and after the learning stage the collective emerging dynamics are studied. In particular, the structured connections *J*_*ij*_, *i, j* = 1, …, *N*, between all neurons are defined through a STDP-based learning rule, forcing the network to learn *P* modular spatiotemporal patterns of spikes **x**^*µ*^(**t**), *µ* = 1, …, *P*, namely:

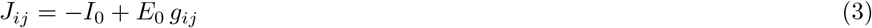

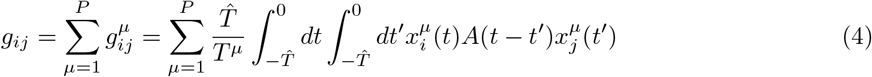

where *I*_0_ is an overall global inhibition and *E*_0_ is a parameter determining the strength of the STDP-based structured term *g*_*ij*_ [22, 25, 30]. The precise timing of spiking activity of neuron *j* in the pattern *µ* is given by 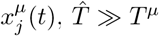 is the learning time, *T* ^*µ*^ is the period of the patterns, the sum over *µ* runs over all the *P* modular patterns to be stored, and *A*(*τ*) is a spike-timing dependent plasticity (STDP) kernel shown in Fig. 1.

Each pattern, acting as teaching signal for cortical plasticity, has a modular structure and it is chosen as a travelling wave on the connectome, involving *G < M* modules. We select the specific sequences of modules of each pattern to be stored in the model, in a way that reflects the structural connectome *W* (*C, C*^*′*^) between cortical regions *C, C*^*′*^, modelled using tractography experimental results. Namely we consider:

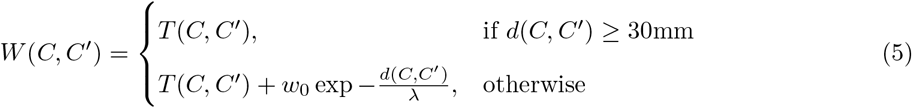

where *T* (*C, C*^*′*^) is the connectivity among cortical ROIs estimated with tractography, i.e. the average number of white tracts averaged over subjects between cortical regions *C* and *C*^*′*^, *w*_0_ is the maximum value in the tractography matrix *T* (*C, C*^*′*^), *d*(*C, C*^*′*^) is the euclidean distance between centroids of regions *C* and *C*^*′*^, and *λ* = 3.33mm is the characteristic length of exponential decay [33, 34]. For anatomically close regions, both myelinated connections from tractography and axonal connections with neighbouring regions (which decay exponentially with distance) are considered, especially since myelinated connections are difficult to measure when fibers are very short.

To extract the sequence of regions involved in pattern *µ*, a closed random walk of G steps is performed through the connectome *W*, using probability *W* (*C, C*^*′*^)*/∑*_*C*′_*W* (*C, C*^*′*^) to go from *C* to *C*^*′*^. Namely, a region *C* is chosen proportionally to ∑_*C′*_ *W* (*C, C*^*′*^) and then a step is done to region *C*^*′*^ according with probabilities *W* (*C, C*^*′*^)*/∑* _*C*′_*W* (*C, C*^*′*^). After *G* + 1 steps, we discard the walk if the (*G* + 1)^*th*^ region does not coincide with the first, until a closed (cyclic) walk is found. This procedure allows to build a travelling wave on the connectome, i.e. a cyclic ordered set of *G* active regions

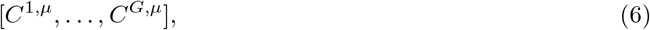

for each pattern *µ* (see Fig. 1).

For each cyclic sequence of regions, a specific periodic sequence of spiking times is extracted for the neurons involved in the pattern *µ* and then this is used in the learning process to build the connectivity *J*_*ij*_. One extracts *K* = *Z/*2 neurons for each of the *G* modules involved in the pattern *µ*, and then we extract *KG* phases in [0 … 2*π*], and assign them to the ordered neurons such that the neurons fires in a sequence respecting the sequence of active modules [*C*^1,*µ*^, …, *C*^*G,µ*^], with partial overlap between modules’ activation. Beginning with the series of neuronal spikes arranged following the prescribed sequence of modules and ensuring no overlap, we execute *n*_*swap*_ = *GK* phase swaps for each pattern. Each swap entails first selecting phases of neurons at a distance *d* with a probability governed by *exp*(*−d*^2^*/*(2*R*^2^)), where *d* is expressed in terms of the number of neurons in the original order, and then swapping the places of the two neurons in the sequence by reassigning their phases. An overlap’s range *R* = 400 is used in this study unless otherwise stated.

In the following, we denote by *C*_*i*_ the module where neuron *i* belongs, and by *C*^*k,µ*^ the *k − th* region/module of the cyclic sequence of pattern *µ*. In essence, we consider *P* modular traveling waves of activity denoted by 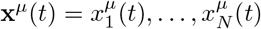, where *µ* = 1 … *P* and:

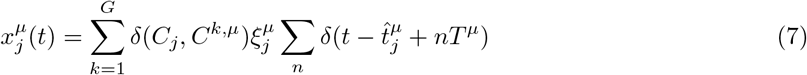

where *C*_*j*_ is the module containing neuron *j*, the Kronecker δ(*C*_*j*_, *C*^*kµ*^) is 1 only if neuron *j* belongs to the *k − th* region active in pattern *µ* and zero otherwise, 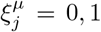 with equal probability, *T* ^*µ*^ is the period of the pattern (here we considered the alpha rhythm using 8 Hz for all patterns to be stored), and 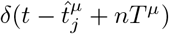is the Dirac delta function, where 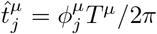is the spike timing of neuron *j* in pattern *µ* and depends on the module *C*_*j*_.

The phase of spiking 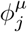 of neuron *j* in pattern *µ* in the active modules *C*^1,*µ*^, .., *C*^*G,µ*^, depends on the module *C*_*j*_ of neuron *j*, in such a way that the neurons fire in a sequence that respects the sequence of active regions [*C*^1,*µ*^, .., *C*^*G,µ*^], with partial overlap between regions, as explained before.

The role of sparse precisely timed patterns of spikes in neural information coding [37] motives the choice of patterns to be stored *x*^*µ*^ as sparse precisely timed spatiotemporal pattern of spiking. In such a way, from equations 4 and 7, we build the microscopic structured connectivity matrix *J*_*ij*_ between the *N* = *ZM* neurons:

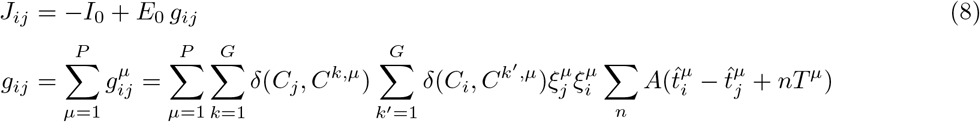

Since each pattern to be stored corresponds to a sequence of modules, extracted using *W* (*C, C*^*′*^) as transition rates, the microscopic *J*_*ij*_ mirrors the modular structure of matrix *W*. The macroscopic connectivity matrix between modules, corresponding to the structured microscopic *g*_*ij*_, is

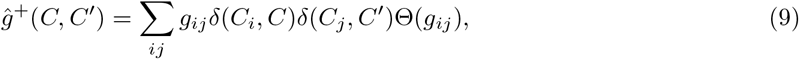

for excitatory connections, and

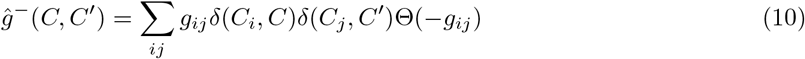

for inhibitory ones, where Θ(*x*) is the Heaviside function, and δ(*C, C*^*′*^) is the Kronecker delta.

The macroscopic connectivity *ĝ*^+^(*C, C*^*′*^) (and *ĝ*^*−*^(*C, C*^*′*^)) between the *M* modules after learning are shown in Fig. 1, togheter with the connectome *W* (*C, C*^*′*^) used to extract stored patterns. The modularity (and the degree of overlap between module’s activity in stored patterns) of the stored patterns is reflected in the degree of modularity (and sparsity) of the connectivity matrix.

To measure the degree of modularity of the microscopic connectivity *g*_*ij*_ w.r.t. the partition into *M* modules, we check whether neurons within the same module exhibit stronger connections among themselves compared to their connections with neurons in other modules. We therefore define a *modularity measure* as

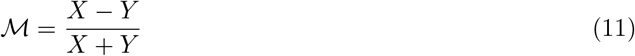

where *X* represents the average strength of internal connections within modules, while *Y* denotes the average strength of connections to neurons external to the module:

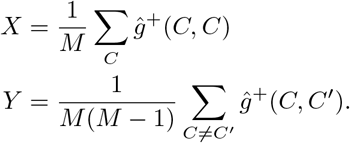

The value of *ℳ* can range in [*−*1, 1]: it is positive if the strength of internal connections is greater than the expected value for a random network, and negative otherwise. In a random network, *ℳ* equals zero on average; conversely, in a network with completely disconnected modules, it equals 1. Here, with *P* = 20 patterns stored using an overlap range of *R* = 400 in a network comprising *N* = 13200 neurons (as used thereafter unless specifically mentioned), we observe *ℳ* = 0.69 for positive connections *ĝ*^+^(*C, C*) and *ℳ* = 0.65 for negative ones *ĝ*^*−*^(*C, C*).

After the learning stage, the *J*_*ij*_ are fixed, and emerging dynamics are studied. It is possible to study the dynamics of the network in two different ways: the spontaneous dynamics in the presence of only noisy term, and the cue-induced dynamics in presence of the external input *I*_*ext*_.

In each pattern, the information is coded both in the binary spatial part, indicating which modules are active and which units inside are active (as in the Hopfield model), and in the temporal part, indicating the precise spiking order (and therefore the precise phase relationships) of the units that are active in that module.

The connections *J*_*ij*_ between neurons are asymmetric because the plasticity depends on the temporal order of the spikings. For this reason, the **attractors of the dynamics** are spatiotemporal patterns with a defined order of spiking of the neurons. The global inhibition reduces the probability that a neuron not participating in the pattern fires incorrectly. The model is similar to the memory storage model of dynamical patterns described in [30], however here we use modular structure, also applying the information *W* (*C, C*^*′*^) as described in the eq. 5 to build connections of the network, and the patterns are not random sequences of neurons.

Notably the network dynamics can selectively and successfully replay one of the stored patterns if a short stimulus resembling the pattern is given. Emergent collective dynamics, reminiscent of one of the stored patterns also occur spontaneously, in the absence of external input.

To measure the success of these replays, the overlap *q*^*µ*^ between the stored pattern *µ* and the network collective activity during the spontaneous dynamics in time interval [*t*_0_, *t*_1_] is measured, in line with [30], defined as:

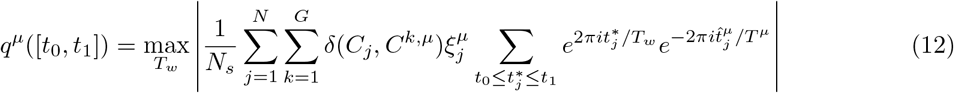

where the sum is taken over spikes *t*^*∗*^ emitted during a specific time interval [*t*_0_, *t*_1_] by neurons belonging to the ones active inside the active modules of the pattern *µ*. Indeed the term 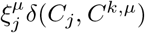 limits the sum to the neurons *j* which actively participate to the stored sequence *µ*. On the contrary *N*_*s*_ represents the total number of observed spikes, including those from neurons not in the pattern, and the maximum is calculated over the time window *T*_*w*_. The overlap tends towards *q*^*µ*^ = 1 when neurons fire in a sequence similar to the original pattern *µ*, even if the replay period *T*_*w*_ is different from the learning stage period *T* ^*µ*^, and neurons silent in the pattern remain inactive. A reduction in replay quality, attributed to non-pattern neurons firing or imprecise replay order, results in a *q*^*µ*^ value less than one. It’s worth noting that the overlap does not assume a specific timescale for the replay, aligning with experimental observations that the reactivation of memory traces during sleep and awake states occurs on a compressed timescale. The order parameter is the largest overlap over all stored patterns:

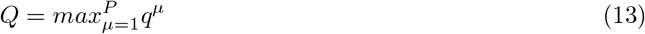

To characterize the phase space, we calculate the fluctuations of the overlap parameter, *χ*_*Q*_ defined as the variance of the order parameter.

### Participants

We enlisted five young adults (4 males, 1 female, average age 29.6 ± 8.87). Each participant was right-handed and native Italian speaker. The criteria for participation included: (1) absence of significant internal, neurological, or psychiatric conditions; and (2) no consumption of substances or medications that might affect MEG/MRI readings. This study adhered to the Helsinki Declaration and received approval from the local Ethics Committee. Written informed consent was obtained from all participants.

### MRI acquisition

Three-dimensional T1-weighted brain scans were captured using a 1.5 Tesla machine (Signa, GE Healthcare) with a 3D Magnetization-Prepared Gradient-Echo BRAVO technique (TR/TE/TI 8.2/3.1/450 ms, 1mm^3^ voxel size, 50% overlapping partitions, 324 sagittal slices encompassing the entire brain). For constructing individual connectomes, diffusion MRI data was gathered using these parameters: Echo-Planar Imaging, TR/TE 12,000/95.5 ms, voxel size of 0.94 × 0.94 × 2.5 mm^3^, 32 diffusion-weighting directions (including 5 B0 volumes). This MRI scanning followed the MEG recording. The diffusion MRI data underwent preprocessing using tools from the FMRIB Software Library (FSL, [http://fsl.fmrib.ox.ac.uk/fsl](http://fsl.fmrib.ox.ac.uk/fsl)). The datasets were adjusted for head movement and eddy current distortions using the *eddy-correct* function, which also reoriented diffusion-sensitizing gradients. A brain mask was created from the B0 images using the Brain Extraction Tool. A diffusion-tensor model was applied to each voxel. Streamlines were produced across the entire brain using deterministic tractography in the Diffusion Toolkit (FACT algorithm, 45° angle threshold, spline-filtering, and using Fractional Anisotropy (FA) maps with a 0.2 threshold for masking). For tractographic analysis, the regions of interest (ROIs) from the AAL atlas and a volumetric version of the DKT ROI atlas, defined in MNI space and masked by the gray matter (GM) probability map from SPM (thresholded at 0.2), were used. FA volumes for each participant were aligned to MNI space using FSL’s FA template and SPM12’s spatial normalization routine. The resulting matrices were inverted and applied to the ROIs to align them with each subject. The quality of this normalization was visually verified. Using custom software in Interactive Data Language (IDL, Harris Geospatial Solutions, Inc), for each subject’s whole-brain tractography and GM ROI set, we calculated the number of streamlines linking each GM ROI pair and their mean tract lengths.

### MEG pre-processing

The preprocessing and source reconstruction of MEG data followed the methods outlined by [35]. The MEG equipment, featuring 163 magnetometers, was developed by the National Research Council of Italy at the Institute of Applied Sciences and Intelligent Systems (ISASI). Briefly, resting-state MEG recordings were split into two segments, each lasting 3 minutes and 30 seconds, with the participants’ eyes closed. The head position was determined using four anatomical reference points and four position coils that were digitized. Additionally, Electrocardiogram (ECG) and Electro-oculogram (EOG) readings were collected.

The MEG signals were initially processed through an anti-aliasing filter, followed by acquisition at a sampling rate of 1024 Hz. These signals were then subjected to a fourth-order Butterworth IIR band-pass filter, operating within the 0.5–48 Hz frequency range. Environmental noise, detected by reference magnetometers, was mitigated using principal component analysis. To eliminate physiological artifacts from the data, such as potential eye blinks and cardiac activity (usually one component), supervised independent component analysis was employed. Noisy channels were identified and removed manually by an expert rater (136 ± 4 sensors were kept).

### Source reconstruction

Neuronal activity time series were reconstructed in 84 regions of interest (ROIs) using the DKT atlas. This was achieved by implementing Nolte’s 2003 volume conduction model and employing the linearly constrained minimum variance (LCMV) beamformer algorithm [59], which was based on the subjects’ own structural MRIs. The source reconstruction was centered on the centroid of each ROI. However, our analysis was narrowed down to 66 ROIs, focusing primarily on cortical areas to ensure the inclusion of the most reliable signals. All preprocessing steps and the source reconstruction process were conducted using the Fieldtrip software [60].

### Fano Factor

To characterize the critical region that opens up where the first-order transition line ends, the Fano Factor of the population spike count [61], and the Coefficient of variation of interspike intervals of modules, are considered. Given spikes counts *n*_*k*_ over time bins *k* = 1 … *n*_bins_ of width δ_bin_, Fano factor is defined as

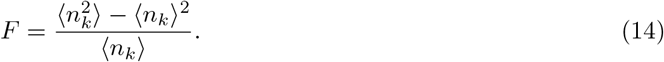

If the event count is extracted from a Poissonian distribution 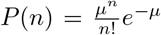, where *μ* = *ρ*(*t*)*δ*_bin_ and *ρ*(*t*) is constant over time, then *F* = 1. If on the other hand, *ρ*(*t*) changes in time, and time bin *δ*_bin_ is larger than the autocorrelation time of *ρ*(*t*), then the Fano is proportional to such autocorrelation time, as discussed in [12]. A long autocorrelation time in *ρ*(*t*) reflects into *F* ≫ 1. It is expected that, in the presence of a critical point the Fano factor value (for enough large time bin *δ*_bin_) exhibits a peak (diverging with the size of the system).

### Avalanche analysis

To investigate the spatial and temporal patterns of transient MEG events, we analyze clusters of significant deflections in artifact-free MEG signals. MEG signals were z-score normalized to have zero mean and unit standard deviation (SD). An avalanche is defined as a continuous time interval in which there is at least one excursion —either positive or negative— beyond threshold in at least one z-scored MEG signal. The aim to detect intervals of large deviations activity is similar to the original works on neuronal avalanches derived from Local Field Potential (LFP) ([38, 62]) and from Multiunit Activity (MUA) spike counts [63]. While for MUA, avalanches were defined as periods of time when the MUA spike count exceeded a positive threshold, and for LFP the excursions were defined for negative potentials due to their close correlation to neuronal spiking, herein, for MEG signals, we considered both positive and negative potentials to identify avalanches in line with previous works on EEG/MEG data ([7, 64–66] A Gaussian distribution of amplitudes is expected to arise from a superposition of many uncorrelated sources. Conversely, MEG amplitude distributions deviate from a Gaussian shape, indicating the presence of spatiotemporal correlations and collective behaviors (Fig. 6). The comparison of the signal distribution to the best Gaussian fit indicates that the two distributions start to deviate from one another around *θ* = *±*3.

An avalanche was thus defined as a series of fluctuations—a continuous period featuring at least one channel exceeding the threshold. We aggregated supra-threshold fluctuations across all 66 signals, defining avalanches as intervals where the cumulative sum was consistently non-zero. Avalanches in this manner are preceded and followed by an empty interval (with sum zero) as usually defined in original avalanches’ works. Size of the avalanche is then defined as the integral of the sum of the absolute values of the signals exceeding the threshold. In the synthetic data, an avalanche is defined as the continuous time interval in which there is at least one module’s rate over zero, where the rate is computed as the number of spikes in 5 ms. The size is defined as the integral of the sum of the rates, and each avalanche is followed and preceded by an empty interval (with firing rate zero).

To evaluate the statistical significance of a power law fit of the observed avalanches’ size distribution we used the *powerlaw* python package [44]. It computes the Kolmogorov-Smirnov statistic based on cumulative distribution functions. For the empirical CDF of data *C*_*emp*_(*s*), and the fitted CDF, *C*_*α*_(*s*), the fit minimizes the KS-distance defined as

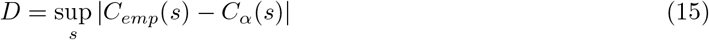

The maximum likelihood estimates of the power law exponents for *P* (*S*) and *P* (*T*) are reported, including the fit error and a systematic error (SE) on the power law fit. The SE is estimated by varying the range of values used for the MLE of the exponents. The power law fit is compared to an exponential fit by evaluating the log-likelihood ratio 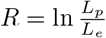 between the likelihood *L*_*p*_ for the power law and *L*_*e*_ for the exponential fit. *R* is positive if the data are more likely to follow a power law distribution, and negative if the data are more likely to follow an exponential distribution. The statistical significance for *R* (p-value) is estimated in the Powerlaw Python package.

### Spearman’s rank correlation coefficient

To quantify the correlation between the empirical degrees of the nodes and the number of times each node initiated a neuronal avalanche in synthetic data, and the correlation between the number of times neuronal avalanches were initiated in a specific brain region in synthetic data and empirical MEG data, we compute Spearman’s rank correlation coefficient (*R*_*S*_). Given two random variables *X* and *Y* and the observations (*x*_*i*_,*y*_*i*_), the Spearman’s rank correlation coefficient is computed as:

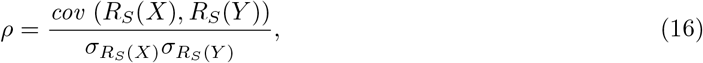

where *R*_*S*_(*X*) is the rank variable associated to *X*, and 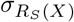 the standard deviation of the rank variable. The value of the rank of a data is its position in the ordered list of the data.

## Data availability statement

The datasets analysed in this study are available from the corresponding author on reasonable request.

## Author contributions

M.A., S.S., P.S., C.C, A.L, L.C. Conceptualization. M.A., S.S., P.S., C.C, A.L, L.C. Methodology. M.A., S.S., P.S., Software. M.A., S.S., P.S., C.C, L.C. Formal analysis. P.S., E.T.L., M.Q., C.G., G.S., S.S. Resources. M.A., S.S., P.S., C.C, L.C. Writing - Original Draft. M.A., S.S., P.S., E.T.L., M.Q., C.G., G.S., V.P., G.M., M.S., S.F., C.C., A.L., L.C. Writing - Review - Editing. M.A., S.S., P.S., V.P., G.M., M.S., S.F., C.C, A.L., L.C. Visualization. S.S., P.S., C.C., L.C. Supervision. S.S., P.S., C.C., L.C. Project administration. S.S., P.S., G.S., S.F. Funding acquisition. All authors read and approved the final manuscript.

## Acknowledgments

M.A., S.F., C.C., A.L., L.C. wish to acknowledge the Italian National Group for Mathematical Physics, GNFM-INdAM. S.F. and C.C. wish to acknowledge ICRANet. Cartoons in Fig. 1, 2 and 6 were created with BioRender.com

